# Resource enrichment and utilization underlie widely observed production and biomass scaling relationships

**DOI:** 10.64898/2026.11.16.767026

**Authors:** Jiayi Song, Oliver Atkinson, David Talmy, Anne Willem Omta

## Abstract

Broadly observed trophic scaling patterns relate the biomass of predators to the biomass of prey, and biomass to productivity. The consistency with which these patterns are observed suggests the existence of governing principles that apply across ecosystems. We present a model for the trophic interaction between phytoplankton and zooplankton that produces the scaling patterns as an emergent property of resource enrichment without invoking the influence of phenotypic or genotypic diversity on system structure. Using formulations with broad empirical support, the model incorporates processes that are common to all ecosystems: resource enrichment, feeding, growth, maintenance, and mortality. Thus, our interpretation of widely observed scaling relationships provides constraints for models that quantify the coupling between ecosystem structure and elemental cycles across terrestrial and aquatic environments.

**Teaser:** A minimal trophic model produces observed macroecological scaling as a result of resource enrichment.

## Introduction

Natural biomes are characterized by diversity at the levels of individual organisms, species, communities, and even entire ecosystems. Even so, there appear to exist general patterns that are observed across different biomes and ecosystems. Paired observations of the biomasses of predators (*Z*) and prey (*P*), and production and biomass, suggest the existence of a general set of power-law scaling relationships (*1, 2*). The predator-prey scaling law was first identified in African savannah ecosystems (*3*) but has since been discovered in systems under very different environmental constraints, including aquatic (*1, 2*) and microbial (*4*) communities. Similarly, the biomass-productivity scaling law is observed across terrestrial (*1*) and aquatic ecosystems (*5, 6*). The ubiquity of these patterns suggests a shared set of underlying governing principles that are yet to be identified.

A recent explanation in the context of phytoplankton and zooplankton communities invoked eco-evolutionary induced changes in phytoplankton size associated with resource enrichment (*7*). However, a challenge arises when this approach is applied to predator-prey pairs that do not exhibit a consistent relationship between biomass and body size, such as mammal carnivores and herbivores. Therefore, it was suggested (*7*) that mammal predator-prey scaling is mediated by a still to be determined prey trait other than size. Here, we construct a trophic model that generates predator-prey and biomass-productivity scaling from resource enrichment, without invoking predator or prey traits. This model is based on the essentials of biological mass transformation and is consistent with several empirical patterns as discussed below.

## Empirical patterns

We focus on the trophic interaction between phytoplankton and zooplankton, because the values of zooplankton feeding parameters are relatively well constrained by a large body of marine observational data (e.g., (*8*)). Further key empirical constraints on the model:

1. Predator-prey scaling (*1*) between the biomasses of zooplankton (*Z*) and phytoplankton (*P*): *Z ∝ P*^*α*^, with *α* the scaling exponent.
2. Paired observations of gross primary production (GPP) and chlorophyll a (Chl-a) indicate a general scaling relationship (*5, 6*): *µ*_*P*_ *P ∝ P*^*β*^, with *µ*_*P*_ the chlorophyll-specific GPP and *β* the scaling exponent.
3. The mean ratio of zooplankton to phytoplankton biomass 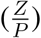 equals 0.1—1.0 (based on (*1*)).
4. Within the naturally occurring range of enrichment, there is stable coexistence between phytoplankton and zooplankton. Although classical trophic models indicate the emergence of limit cycles under nutrient enrichment (*9, 10*), such behavior is generally not observed in natural plankton communities (*11*).

We estimated the scaling exponents *α* and *β* from published paired observations of phytoplankton and zooplankton biomass and Chl-a and GPP, respectively (Fig. 1). We used Reduced Major Axis (or Type 2) regression, because there likely exists error variance in both *Z* and *P* (*13*). For this same reason, we used as a measure of the goodness of fit the orthogonal *R*^2^ (indicated as 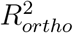). This 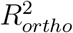 is equivalent to the proportion of variance explained by the first component in Principal Component Analysis (*14*). For a fair comparison, we also used Type 2 regression when estimating scaling exponents from model output.

**Figure 1:**
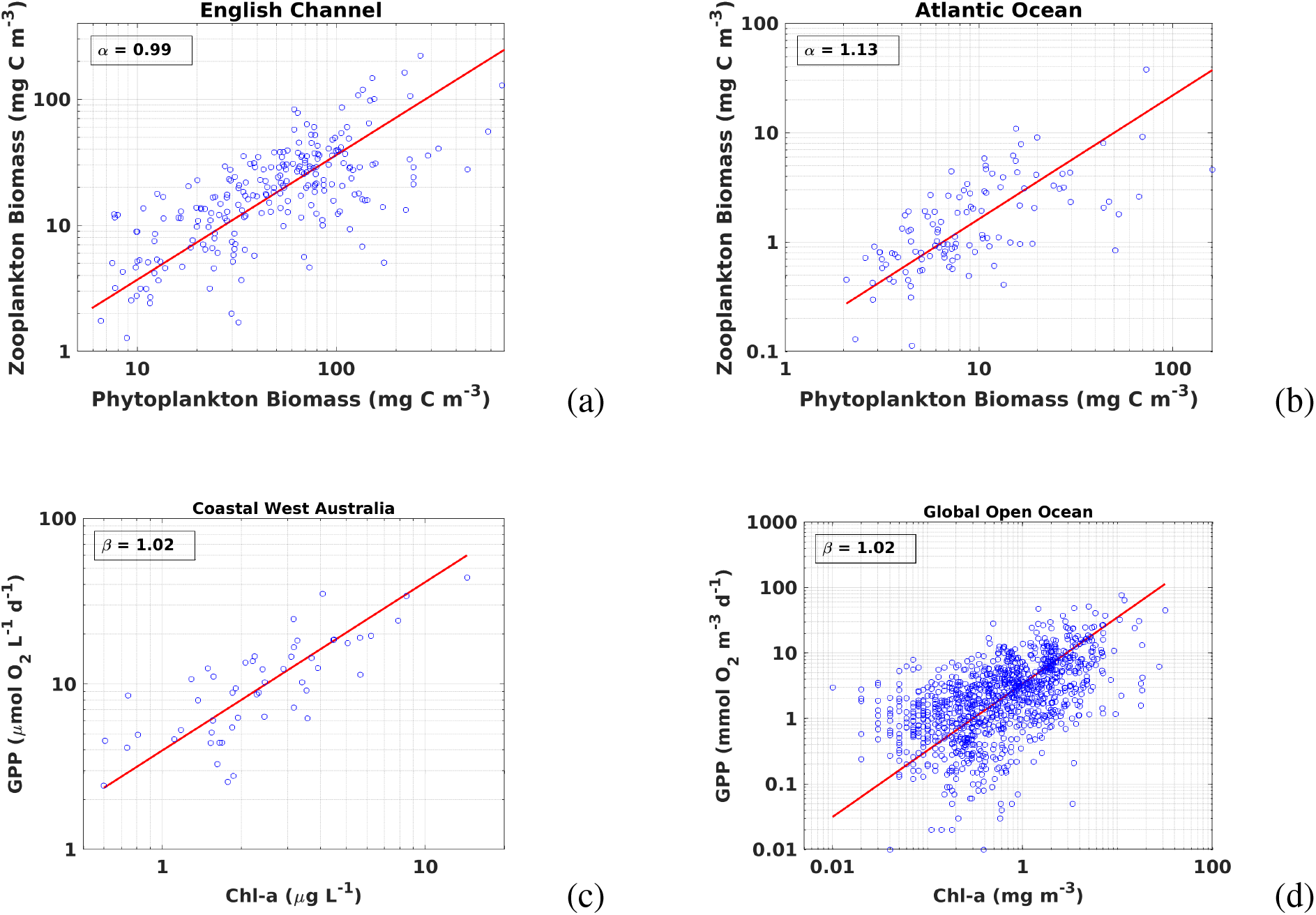
Observed phytoplankton vs zooplankton biomass (blue dots) with power-law regression lines (red) on log-log scale to estimate the scaling exponent *α* (a, b). Observed gross primary production vs chlorophyll (blue dots) with power-law regression lines (red) on log-log scale to estimate the scaling exponent *β* (c, d). a: English Channel (*12*), 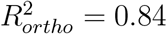 b: Atlantic Ocean (*12*), 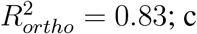 Coastal West Australia (*6*),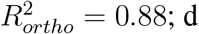 : Global Open Ocean (*5*),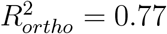.

We note that, while Chl-a is a widely used proxy for biomass, there is variation in the chlorophyll-to-carbon ratio (Chl:C) associated with taxonomy (*15,16*) and photoacclimation (*17– 19*). For our analysis, a potentially confounding effect is that the Chl:C ratio tends to increase with nutrient enrichment (*20, 21*). In particular, this could lead to an underestimation of the value of the scaling exponent *β*. That said, the range in the Chl:C ratio of phytoplankton across different marine communities is estimated to be around a factor 10 (*22, 23*), which is about 2 orders of magnitude smaller than the range of Chl-a in the data under consideration. In other words, most of the observed variation in Chl-a reflects a variation in phytoplankton biomass.

## Model construction

Our nutrient-phytoplankton-zooplankton (NPZ) model is schematically depicted in Fig. 2. It represents the processes of phytoplankton growth and nutrient uptake, zooplankton grazing and mortality, and phytoplankton and zooplankton maintenance through the following equations:

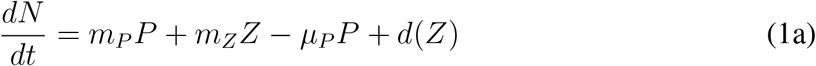

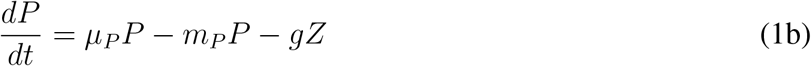

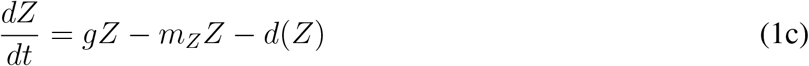

**Figure 2:**
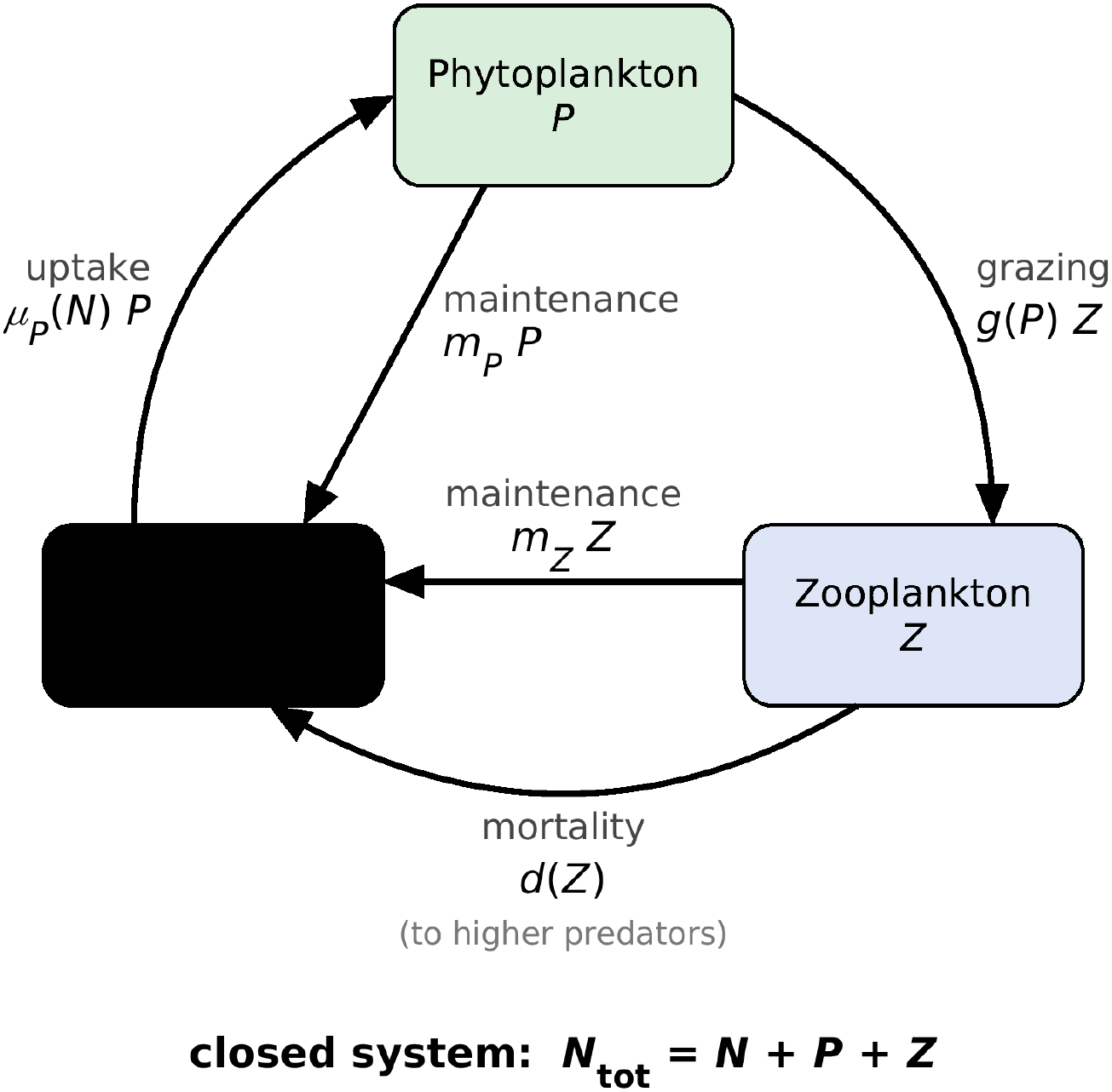
Schematic depiction of the model with the different pools (nutrient, phytoplankton, zooplankton) and trophic interactions (indicated by arrows).

In the set of equations (1), *N* is the nutrient concentration, while *P* and *Z* are the phyto-plankton biomass and zooplankton biomass (all these three variables are in units of *µ*M N). The maintenance loss rates of phytoplankton and zooplankton biomass are represented by *m*_*P*_ and *m*_*Z*_, respectively. We used saturating functions to describe all the feeding interactions, since such functions are ubiquitous in mass transformations in biological systems across scales (*24*). In all simulations, we used a Holling Type 2 formulation for the phytoplankton nutrient uptake rate 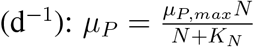 (with maximum uptake rate *µ*_*P,max*_ and half-saturation constant *K*_*N*_). Losses due to phyto- and zooplankton maintenance and zooplankton mortality are recycled into the nutrient pool, creating a closed system. Furthermore, we used a hyperbolic formulation for the zooplankton mortality: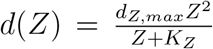, with *d*_*Z,max*_ a maximum mortality rate and *K*_*Z*_ a half-saturation constant for predation on zooplankton. The hyperbolic mortality formulation is based on a combination of Holling Type 2 predation on zooplankton by higher trophic levels (e.g., macrozooplankton, small fish) and a linear proportionality between the density of zooplankton and its predator (*25*). A key consideration for choosing this formulation is the following. We estimated *β ≈* 1 from the empirical data (*5, 6*), which means that the phytoplankton productivity vs biomass scaling relationship is approximately linear: *µ*_*P*_ *P ∝ P* . Based on equation (1a), *µ*_*P*_ *P* = *m*_*P*_ *P* + *m*_*Z*_*Z* + *d*(*Z*) at steady state. Since *P* and *Z* must be approximately linearly related (to be consistent with our estimated *α ≈* 1) and *m*_*P*_ *P* and *m*_*Z*_*Z* are linear functions of *P* and *Z*, the zooplankton mortality *d*(*Z*) needs to be linear in *Z* (and thus *P*) for the productivity *µ*_*P*_ *P* to be linear in *P* at steady state. However, a hyperbolic zooplankton mortality gives a more realistic response to nutrient enrichment than a linear zooplankton mortality (*25*). By taking *K*_*Z*_ = 0.1 *µ*M N (instead of *K*_*Z*_ = 1.0 *µ*M N as in a previous study (*25*)), the model approaches a linear mortality with the hyperbolic formulation. The values of the fixed parameters *µ*_*P,max*_, *K*_*N*_, *K*_*Z*_, *m*_*P*_, and *m*_*Z*_ are listed in Table S1.

It is unclear which commonly used grazing formulation (Holling Type 2, Holling Type 3, or Ivlev) is the most consistent with experimental feeding curves (*26*). Furthermore, it is questionable how well these experimentally determined feeding curves extrapolate to the natural environment (*27*). Therefore, we performed simulations and analyses with several grazing formulations commonly used in marine biogeochemical models (*28*):

- Holling Type 2 (with feeding threshold):

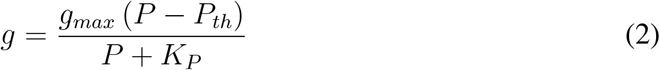

- Holling Type 3:

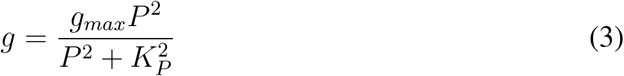

- Ivlev:

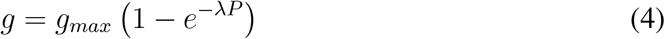

In eqs. (2–4), *g*_*max*_ is the maximum grazing rate, *K*_*P*_ is a half-saturation constant for grazing, *P*_*th*_ is a feeding threshold, and *λ* is a capture efficiency parameter; values for these parameters are listed in Table S2.

## Results

We used the bifurcation analysis package Auto (*29*) to analyze the modeled scaling behavior along a eutrophication gradient imposed by varying the total nutrient inventory *N*_*tot*_ *≡ N* + *P* + *Z* between 0.2 and 10 *µ*M N, which approximately covers the observed global range in total surface nitrogen across oceans and lakes based on biomass (*1, 30*). In cases where the zooplankton goes extinct at *N*_*tot*_ = 0.2 *µ*M N, the minimum value of *N*_*tot*_ used was equal to the minimum concentration at which the zooplankton can establish itself: *N*_*tot*_ = 0.28 *µ*M N for Holling Type 3 grazing with parameters from the World Ocean Model of Biogeochemistry and Trophic dynamics (WOMBAT) (*31*) and *N*_*tot*_ = 0.35 *µ*M N for Holling Type 3 grazing with parameters from the Ocean Ecosystem Component model, version 2 (OECO-v2) (*32*).

Since the zooplankton mortality rate is uncertain (*25*), we used constraint (1) to tune *d*_*Z,max*_. More specifically, we varied *d*_*Z,max*_ until *α ≈* 1 for each grazing formulation. Then, we evaluated whether the results (summarized in Table 1) are consistent with constraints (2–4). Holling Type 2 grazing with parameters from the isopycnic coordinate HAMburg Ocean Carbon Cycle (iHAMOCC) model (*33*) and the Marine Biogeochemistry Library (MARBL) (*34*), Holling Type 3 grazing with parameters from WOMBAT, and Ivlev grazing with parameters from the Canadian Ocean Ecosystem (CanOE) model (*36*) all gave *β ≈* 1 (consistent with constraint 2).

**Table 1:**
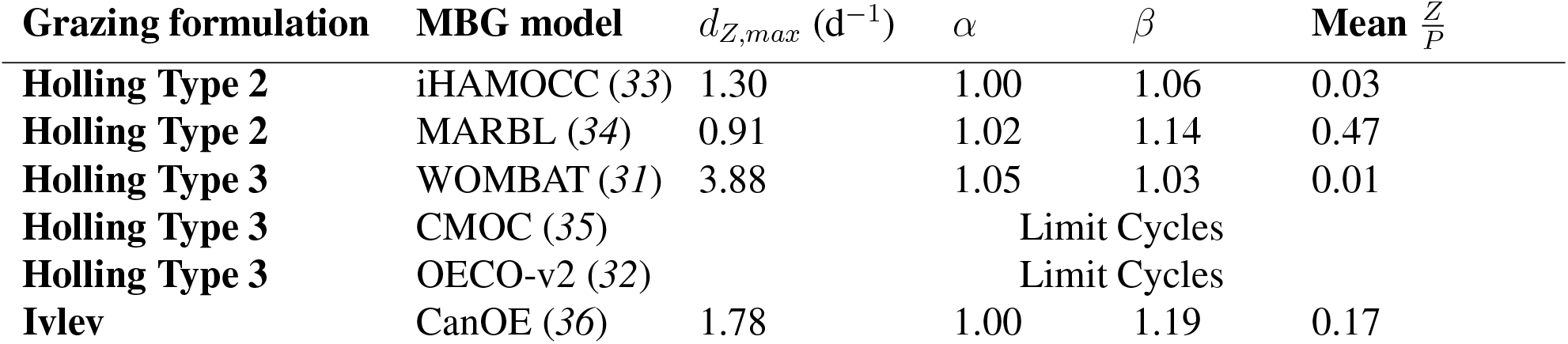
Analysis results for different grazing formulations, with the marine biogeochemical (MBG) model from which the grazing parameter values were taken, the max. zooplankton mortality (*d*_*Z,max*_) needed to obtain *α* ≈ 1, the scaling exponents *α* and *β*, and the mean 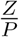 ratio

| **Grazing formulation** | **MBG model** | *d*<sub>*Z,max*</sub> (d<sup>−1</sup>) | *α* | *β* | **Mean** $\frac{Z}{P}$ |
|---|---|---|---|---|---|
| **Holling Type 2** | iHAMOCC (*33*) | 1.30 | 1.00 | 1.06 | 0.03 |
| **Holling Type 2** | MARBL (*34*) | 0.91 | 1.02 | 1.14 | 0.47 |
| **Holling Type 3** | WOMBAT (*31*) | 3.88 | 1.05 | 1.03 | 0.01 |
| **Holling Type 3** | CMOC (*35*) | Limit Cycles |  |  |  |
| **Holling Type 3** | OECO-v2 (*32*) | Limit Cycles |  |  |  |
| **Ivlev** | CanOE (*36*) | 1.78 | 1.00 | 1.19 | 0.17 |

However, the mean 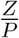 ratio was well below 0.1 for the iHAMOCC and WOMBAT grazing, which is inconsistent with constraint 3. Furthermore, the mean 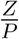 ratio was somewhat low for the CanOE grazing. As we were tuning *d*_*Z,max*_ to approach *α* = 1, we found that Holling Type 3 grazing with parameters from the Canadian Model of Ocean Carbon (CMOC) (*35*) and OECO-v2 gave limit cycles toward higher values of *N*_*tot*_ (inconsistent with constraint 4). Overall, the results from the model with the MARBL Holling Type 2 grazing parameterization are the most consistent with all our constraints. Indeed, the simulated relationships between zooplankton and phytoplankton biomass and between phytoplankton productivity and phyto-plankton biomass were close to linear (Fig. 3), the mean 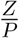 ratio was equal to 0.47, and the system was stable throughout the range of *N*_*tot*_ under consideration.

**Figure 3:**
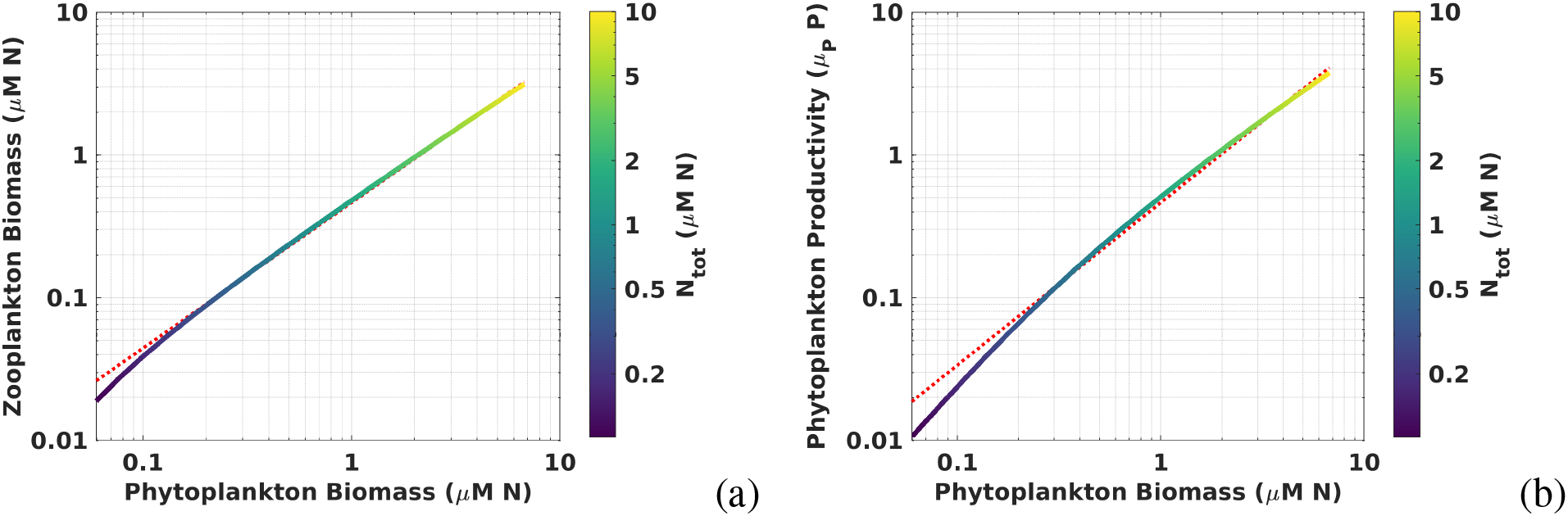
The model with the MARBL (*34*) grazing parameterization and *d*_*Z,max*_ = 0.91 d^*−*1^ yields *α* = 1.02, *β* = 1.14, which is roughly consistent with the values from the empirical data. The curves indicate the simulated zooplankton biomass vs phytoplankton biomass (a) and phytoplankton productivity vs phytoplankton biomass (b), with the different colors corresponding to eutrophication levels. Exactly linear scaling (*α* = 1, *β* = 1) is indicated by the red lines.

We investigated the stability of the model with MARBL grazing in more detail by exploring the impact of nutrient enrichment (increasing *N*_*tot*_) for different values of *d*_*Z,max*_, *g*_*max*_, and *K*_*P*_ . We focused on these three parameters because of the uncertainty in *d*_*Z,max*_ (*25*) and the large observed range in *g*_*max*_ and *K*_*P*_ (*8*). Examples of stable zooplankton-phytoplankton coexistence (consistent with criterion 4) and limit cycles (inconsistent with criterion 4) are provided in the Supplementary Figs. S1a and S1b, respectively. Limit cycles occurred toward high values of *N*_*tot*_ combined with low values of *d*_*Z,max*_ (Fig. 4a), moderately high values of *g*_*max*_ (Fig. 4b), or low values of *K*_*P*_ (Fig. 4c), all of which lead to relatively high zooplankton abundances and thus a top-heavy trophic chain. Such top-heaviness determining the occurrence of limit cycles is consistent with findings from a previous study (*25*). For the current model, the Hopf bifurcation corresponded to a 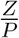 ratio of around 1. In addition, the stability of the system appeared to be somewhat affected by nutrient enrichment. That is, the Hopf bifurcation occurred at a 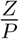 ratio of about 1.5 for *N*_*tot*_ = 0.6 *µ*M N and at a 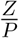 ratio of about 0.8 for *N*_*tot*_ = 10 *µ*M N. Finally, stable behavior under eutrophication also occurred for very high values of *g*_*max*_ or very low values of *K*_*P*_ . This appears to be due to a strong top-down control by the zooplankton. The resulting steady state is characterized by high 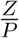 ratios, low concentrations of both phyto- and zooplankton, and a low nutrient utilization (i.e., 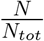 close to 1). Overall, realistic 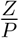 ratios (0.1—1.0) were associated with stable coexistence between the phytoplankton and zooplankton populations. The large range of parameter combinations with stable coexistence also ensures that the system would not easily be pushed into limit cycles due to noise and variability in the parameters.

**Figure 4:**
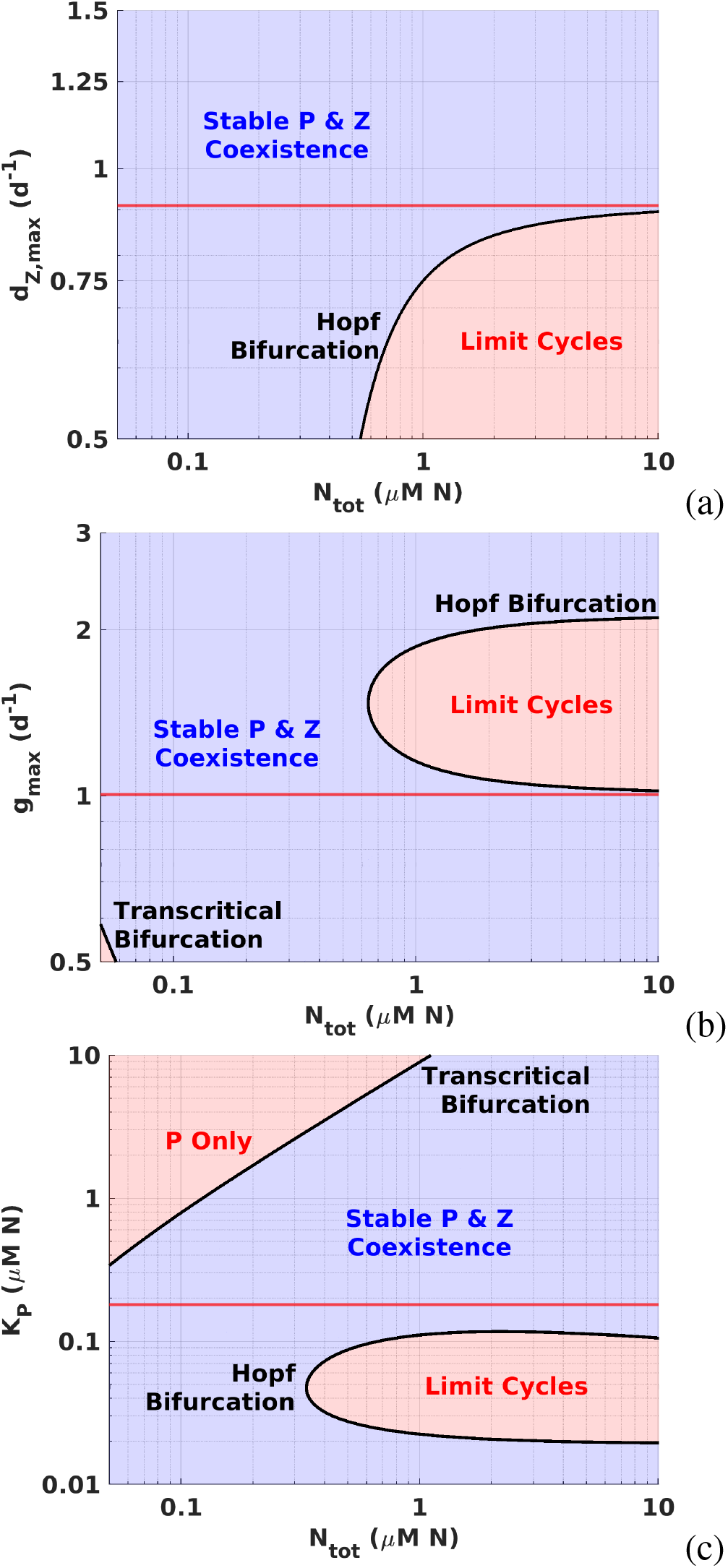
The model with MARBL grazing exhibits stable coexistence of phytoplankton and zooplankton upon enrichment for most of the ranges of *d*_*Z,max*_, *g*_*max*_, and *K*_*P*_, as indicated by the bifurcation diagrams: *(a) N*_*tot*_ and *d*_*Z,max*_ varied, fixed *g*_*max*_ = 1.00 d^*−*1^, *K*_*P*_ = 1.5*10^*−*4^ *µ*M N; *(b) N*_*tot*_ and *g*_*max*_ varied, fixed *d*_*Z,max*_ = 0.91 d^*−*1^, *K*_*P*_ = 1.5*10^*−*4^ *µ*M N; *(c) N*_*tot*_ and *K*_*P*_ varied, fixed *d*_*Z,max*_ = 0.91 d^*−*1^, *g*_*max*_ = 1.00 d^*−*1^.

## Discussion

We have constructed a model for the trophic interaction between phytoplankton and zooplankton that gives results consistent with both the trophic scaling relationships for predators and prey (*1, 2*) and phytoplankton productivity (*5, 6*) as the nutrient inventory is changed. Furthermore, the model produced reasonable zooplankton-to-phytoplankton biomass ratios, as well as stable coexistence between phytoplankton and zooplankton in a large range of parameter space. Our model requires a minimal set of assumptions with broad empirical support. Most importantly, the growth rates of predators and prey follow saturating curves with respect to their respective resources (prey for predators; nutrients for prey). This simple principle is encapsulated in the Holling Type 2 functional response, which is widely observed (*37*), and leads to model predictions consistent with empirical macroecological scaling relationships (*1, 5, 13*), the observed ratio between zooplankton and phytoplankton biomass, and the response to nutrient enrichment. Previous studies have found unstable population dynamics to be rather prevalent with Holling Type 2 grazing (*8, 38, 39*), which can lead to a poor fit to local seasonal dynamics (*40*). Our results suggest that this unstable behavior is primarily the result of a top-heavy trophic chain, which in turn depends on the choice of grazing and zooplankton mortality parameter values. Although predator-to-prey biomass ratios higher than 1 are rare in natural systems (*1, 2*), several examples of such inverted trophic chains have been reported (*41, 42*). It has been hypothesized that stable top-heavy trophic chains may be maintained through exogenous subsidies such as inward migration of predators orprey (*43*). Our results suggest that a top-heavy trophic chain can also be stabilized through high per-capita predation rates (as achieved by high values of *g*_*max*_ or low values of *K*_*P*_ in our model).

Prior discussions surrounding biomass and productivity scaling patterns have centered around a set of 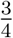 power laws. The 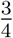 power law implies that the ratio of predators to prey decreases as systems become more resource rich. This ‘sublinear’ behavior was established with a type of model fitting that assumes error-free sampling of the prey biomass (Ordinary Least Squares, Type 1 regression). Such an assumption may be appropriate for controlled laboratory experiments where an independent variable can be reliably measured, but it may not be applicable in ecological contexts where unresolved processes introduce error (*44*). There is no reason to assume that these unresolved processes would, in relative terms, have a stronger or a weaker effect on predator than on prey biomass. In such a case, Type 2 (e.g., Reduced Major Axis) regression provides a relatively unbiased estimate of the slope of the relationship, since it represents the geometric mean of the values obtained when assuming that all the error is in one variable or the other variable. For both the observational data and the model results, we have therefore used Type 2 regression. This yields approximately linear scaling between predators and prey and invariant predator-to-prey ratios as biomass increases (*13*), which we aimed to reproduce with our model.

Recently, an agent-based model without traits generated power-law scaling between average predator and prey densities in the regime with predator-prey cycles (*45, 46*). Furthermore, a modified Lotka-Volterra model reproduced predator-prey scaling with populations at steady state (*47*). However, these models did not include an explicit nutrient or resource limiting the prey population growth. As a result, the representation of resource enrichment was not empirically grounded. Furthermore, these models aimed to reproduce the 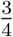 power-law scaling behavior that emerged under the assumption of an error-free independent variable. As discussed above, we aimed to reproduce the approximately linear scaling consistent with error variance in both variables.

Real-world aquatic ecosystems are significantly more complex than the nutrient-phytoplankton-zooplankton model we constructed, which aggregates diverse communities of predators and prey into single groups with fixed properties. Both phyto- and zooplankton consist of many species occupying different ecological niches living in a dynamic ocean environment. As a result, the population dynamics could be affected by spatial heterogeneity and the structure of the food web. However, a recent study using a three-dimensional ocean model found that the biomass scaling is affected more by the grazing and zooplankton mortality formulations than by the food-web structure (*48*). A related consideration is our omission of zooplankton ‘active switching’, i.e., preferable targeting of more abundant prey species (*49*). Although active switching is not a universally observed behavior, its inclusion in ecosystem models tends to enhance the stability and species diversity of the modeled food web (*50*). Even so, models including active switching by zooplankton typically conform to Holling Type 2 or 3 for grazing on the aggregate phytoplankton community (*50*). Therefore, we hypothesize that the insights from our highly idealized study apply to these more complex systems if the aggregate phytoplankton and zooplankton communities (rather than individual species or groups) are considered. In future work, this hypothesis could be tested through simulations with models of increasing complexity.

Although we focused on phytoplankton and zooplankton, our findings likely have a broader applicability across different ecosystems and trophic levels. The Holling Type 2 saturating functional response is ubiquitous in trophic interactions (*24*). Moreover, our model resolves the basic processes of feeding, growth, maintenance, and mortality, which occur in any predatorprey pair. In other words, the broadly observed macroecological patterns reproduced by our model can be a general consequence of trophic interactions.

## Acknowledgments

The authors would like to thank E. Carr and G. Britten for helpful discussions and suggestions.

## Authors contributions

J.S. performed conceptualization, investigation, analyses, and writing – original draft, O.A. performed investigation, analyses, and writing – review & editing, D.T. performed conceptualization and writing – review & editing, A.W.O. performed conceptualization and writing – review & editing.

## Data, code, and materials availability

This study did not generate any primary data. The code used in the simulations and analyses is available through A.W.O.’s GitHub page: https://github.com/AWO-code/NPZ.

## Supplementary materials

Tables S1 and S2

Figure S1

**Table S1:** Fixed parameters with interpretations, values, and units.

| Parameter | Description | Value | Unit |
|---|---|---|---|
| *μ*<sub>*P,max*</sub> | Maximum phytoplankton growth rate | 1.0 | d<sup>−1</sup> |
| *K*<sub>*N*</sub> | Half-saturation constant for N uptake | 0.1 | *μ*M N |
| *K*<sub>*Z*</sub> | Half-saturation constant for predation on zooplankton | 0.1 | *μ*M N |
| *m*<sub>*P*</sub> | Phytoplankton maintenance loss | 0.1 | d<sup>−1</sup> |
| *m*<sub>*Z*</sub> | Zooplankton maintenance loss | 0.1 | d<sup>−1</sup> |

**Table S2:** Grazing formulations used with parameter values and the marine biogeochemical (MBG) model in which these parameter values are used. These were taken from a compilation of published marine biogeochemical models (*28*) (using a Redfield C:N ratio of 6.6 for conversion from *mu*M C to *µ*M N).

| **Grazing formulation** | *g*<sub>*max*</sub> (d<sup>−1</sup>) | *K*<sub>*P*</sub> (*µ*M N) | *P*<sub>*th*</sub> (*µ*M N) | *λ* (*µ*M N) | **MBG model** |
|---|---|---|---|---|---|
| **Holling Type 2** | 1.20 | 1.48 | 1.5*10<sup>−5</sup> | NA | iHAMOCC (*33*) |
| **Holling Type 2** | 1.00 | 0.18 | 1.5*10<sup>−4</sup> | NA | MARBL (*34*) |
| **Holling Type 3** | 1.58 | 1.00 | NA | NA | WOMBAT (*31*) |
| **Holling Type 3** | 2.00 | 0.20 | NA | NA | CMOC (*35*) |
| **Holling Type 3** | 2.00 | 1.42 | NA | NA | OECO-v2 (*32*) |
| **Ivlev** | 1.70 | NA | NA | 1.65 | CanOE (*36*) |

**Figure S1:**
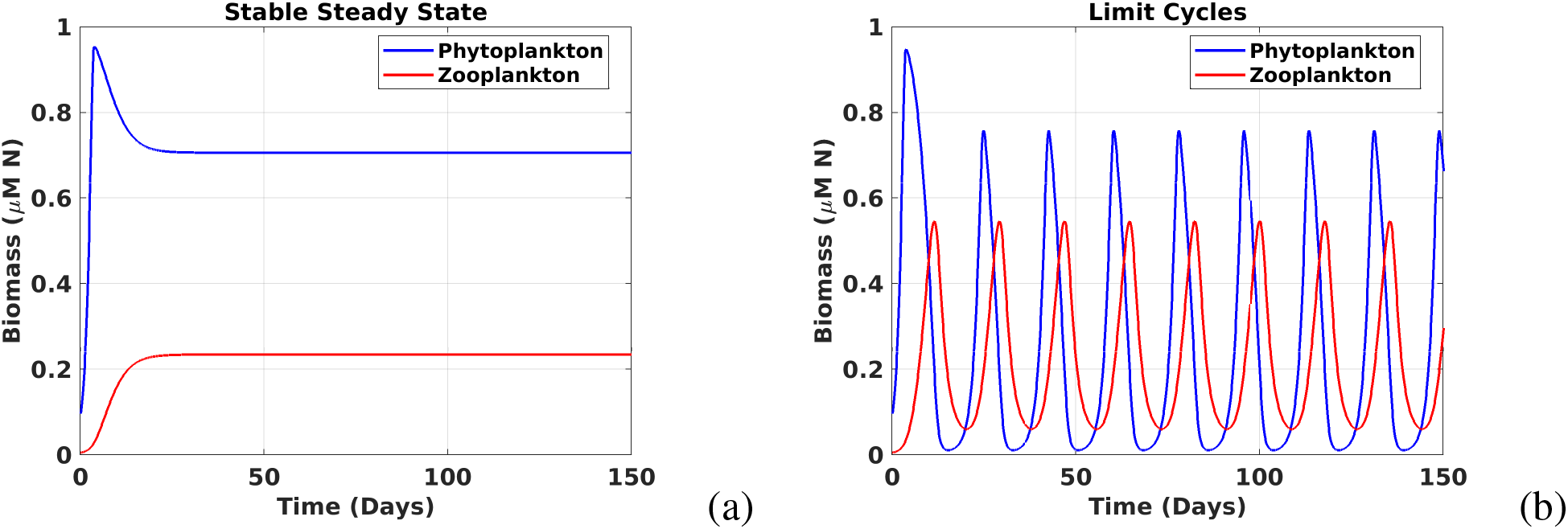
Examples of phytoplankton (blue) and zooplankton (red) biomass vs. time for (a) stable coexistence and (b) limit cycles. The simulations were performed using the MARBL (*34*) grazing parameterization for *N*_*tot*_=1.0 *µ*M and *d*_*Z,max*_ = 1.0 d^*−*1^ (a), *d*_*Z,max*_ = 0.5 d^*−*1^ (b).

