## Supplementary Tables S1 and S2, Figure S1 for "Resource enrichment and utilization underlie widely observed production and biomass scaling relationships"

### Supplementary materials

Tables S1 and S2

Figure S1

| Parameter | Description | Value | Unit |
|---|---|---|---|
| $\mu_{P,max}$ | Maximum phytoplankton growth rate | 1.0 | d$^{-1}$ |
| $K_N$ | Half-saturation constant for N uptake | 0.1 | $\mu$M N |
| $K_Z$ | Half-saturation constant for predation on zooplankton | 0.1 | $\mu$M N |
| $m_P$ | Phytoplankton maintenance loss | 0.1 | d$^{-1}$ |
| $m_Z$ | Zooplankton maintenance loss | 0.1 | d$^{-1}$ |

Table S1: Fixed parameters with interpretations, values, and units

| Grazing formulation | $g_{max}$ (d$^{-1}$) | $K_P$ ($\mu$M N) | $P_{th}$ ($\mu$M N) | $\lambda$ ($\mu$M N) | MBG model |
|---|---|---|---|---|---|
| **Holling Type 2** | 1.20 | 1.48 | 1.5*10$^{-5}$ | NA | iHAMOCC (*1*) |
| **Holling Type 2** | 1.00 | 0.18 | 1.5*10$^{-4}$ | NA | MARBL (*2*) |
| **Holling Type 3** | 1.58 | 1.00 | NA | NA | WOMBAT (*3*) |
| **Holling Type 3** | 2.00 | 0.20 | NA | NA | CMOC (*4*) |
| **Holling Type 3** | 2.00 | 1.42 | NA | NA | OECO-v2 (*5*) |
| **Ivlev** | 1.70 | NA | NA | 1.65 | CanOE (*6*) |

Table S2: Grazing formulations used with parameter values and the marine biogeochemical (MBG) model in which these parameter values are used. These were taken from a compilation of published marine biogeochemical models (*7*) (using a Redfield C:N ratio of 6.6 for conversion from $mu$M C to $\mu$M N).

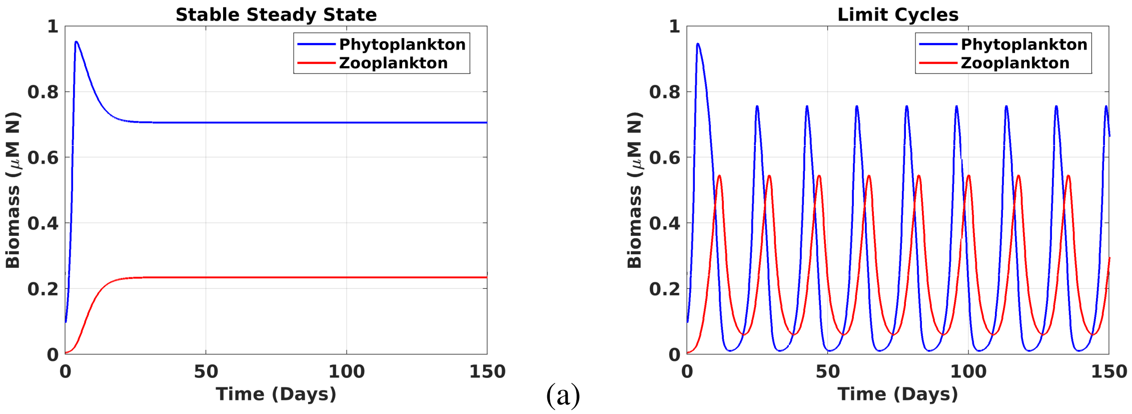

(a) (b)

Figure S1: Examples of phytoplankton (blue) and zooplankton (red) biomass vs. time for (a) stable coexistence and (b) limit cycles. The simulations were performed using the MARBL (*2*) grazing parameterization for $N_{tot}$=1.0 $\mu$M and $d_{Z,max} = 1.0$ d$^{-1}$ (a), $d_{Z,max} = 0.5$ d$^{-1}$ (b).


3. R. M. Law, *et al.*, The carbon cycle in the Australian Community Climate and Earth System Simulator (ACCESS-ESM1) – Part 1: Model description and pre-industrial simulation. *Geoscientific Model Development* **10**, 2567–2590 (2017).

4. K. Zahariev, J. R. Christian, K. L. Denman, Preindustrial, historical, and fertilization simulations using a global ocean carbon model with new parameterizations of iron limitation, calcification, and $N_2$ fixation. *Progress in Oceanography* **77**, 56–82 (2008).

5. T. Hajima, *et al.*, Development of the MIROC-ES2L Earth system model and the evaluation of biogeochemical processes and feedbacks. *Geoscientific Model Development* **13**, 2197–2244 (2020).

6. J. R. Christian, *et al.*, Ocean biogeochemistry in the Canadian Earth System Model version 5.0.3: CanESM5 and CanESM5-CanOE. *Geoscientific Model Development* **15**, 4393–4424 (2022).

7. T. Rohr, A. J. Richardson, A. Lenton, M. A. Chamberlain, E. H. Shadwick, Zooplankton grazing is the largest source of uncertainty for marine carbon cycling in CMIP6 models.

*Communications Earth & Environment* **4**, 212 (2023).
